# Redox-dependent binding and conformational equilibria govern the fluorescence decay of NAD(P)H in living cells

**DOI:** 10.1101/2024.12.13.628382

**Authors:** Thomas S. Blacker, Nimit Mistry, Nicoletta Plotegher, Elizabeth R. Westbrook, Michael D. E. Sewell, John Carroll, Gyorgy Szabadkai, Angus J. Bain, Michael R. Duchen

## Abstract

When probed using fluorescence lifetime imaging microscopy (FLIM), the emission from reduced nicotinamide adenine dinucleotide (NADH) and its phosphorylated form NADPH have shown promise as sensitive intrinsic reporters of metabolism in living systems. However, an incomplete understanding of the biochemical processes controlling their fluorescence decay makes it difficult to draw unambiguous conclusions from NAD(P)H FLIM data. Here we utilised time-resolved fluorescence anisotropy imaging to identify multiple enzyme binding configurations in live cells associated with lifetimes both longer and shorter than unbound NAD(P)H. FLIM, combined with mathematical and computational modelling, revealed that the redox states of the NAD and NADP pools control the steady-state equilibrium of binding configurations, which in-turn determines the observed fluorescence decay. This knowledge will be foundational to developing the accurate interpretation of NAD(P)H FLIM.

## Introduction

The pools of nicotinamide adenine dinucleotide (NAD) and its phosphorylated analogue NADP play essential roles in cellular metabolism, ferrying electrons to and from redox reactions crucial to energy production, biosynthesis, antioxidant defence and cellular homeostasis^1^. Preservation of both cofactors is linked to healthy ageing^2,3^, and their disruption is implicated in various diseases^4^, making them promising targets for therapeutic intervention^5^. In their reduced (electron carrying) forms, NADH and NADPH are intrinsically fluorescent, and this is lost upon oxidation to NAD+ and NADP+. As the fluorescence spectra of the two reduced cofactors are identical, the combined emission from living tissues is often labelled NAD(P)H^6^. These characteristics have been utilised for the non-invasive interrogation of cellular metabolism since the 1950’s^7^. Early experiments monitored NAD(P)H intensity using spectrofluorometry to address fundamental questions in respiratory chain activity and tissue oxygenation^8,9^. The development of laser scanning confocal microscopy enabled subcellular imaging of NAD(P)H^10^, providing insights into the role of mitochondrial dysfunction in disease^11^. These intensity-based techniques continue to be applied on modern confocal and two-photon microscopes^12,13^. However, there is considerable interest in the extra dimensions of metabolic information that may be obtained from living systems through technologies that exploit the quantitative precision of time-resolved NAD(P)H fluorescence^14,15^.

The average duration of excited state occupation following optical absorption defines the fluorescence lifetime of a molecule. This is highly sensitive to its local environment and interactions, motivating the application of fluorescence lifetime imaging microscopy (FLIM) of NAD(P)H as a label-free metabolic probe^6^. In live cells, NAD(P)H FLIM typically resolves two lifetimes at each pixel: a short component denoted *τ*_1_(300-500ps) associated with freely diffusing species, and a longer component *τ*_2_ (1500-4500ps) attributed to enzyme-bound forms^15,16^. The relative abundance of the two species is also quantified, typically reported as the percentage of the emitting population exhibiting the longer lifetime, *α*_2_. Differences in these parameters have been described in a range of disease models over the last two decades^14^, leading to the development of diagnostic instrumentation^17^. Despite these advances, NAD(P)H FLIM has yet to be established as a routine technique for detailed metabolic characterisation as interpretation in terms of underlying cellular biochemical processes remains poorly understood. We address these issues here.

We have previously reported that the fluorescence lifetimes of NADH and NADPH are differentially altered by various binding configurations to oxidoreductase enzymes in solution^18^. In the present study, we have investigated the role that these configurations play in NAD(P)H FLIM measurements performed in living cells. By combining time-resolved fluorescence anisotropy imaging (trFAIM) with FLIM measurements in a range of living cell models under different conditions, mathematical modelling of redox-dependent binding equilibria, and computational modelling of the decay fitting process, we have found that the cellular [NAD+]:[NADH] and [NADP+]:[NADPH] redox balances control steady-state equilibria of binding configurations for each cofactor, dictating the lifetime distribution of fluorescent species present. This results in a fluorescence decay with at least seven components, which approximates to a biexponential under the limited signal-to-noise conditions when imaging living samples. We then show that the lifetimes of the two components (*τ*_1_and *τ*_2_) and their relative weighting (*α*_2_, see methods) are each sensitive to the redox states of the NAD or NADP pools. Understanding the biomolecular processes that influence NAD(P)H fluorescence lifetime variations will pave the way towards the meaningful interpretation of FLIM experiments in terms of underlying cell metabolism, applied to both fundamental biology and clinical diagnostic measurements.

## Results

### Enzyme bound species with fluorescence lifetimes both longer and shorter than free NAD(P)H are present in living cells

NAD and NADP impart their cellular functions through oxidoreductases, the catalytic process of which involves conformational changes in both enzyme and cofactor^19–21^. In the open conformation of the enzyme, NAD(P) binds at its adenine end, leaving a mobile nicotinamide moiety. In the closed conformation, typically promoted by substrate binding, the nicotinamide becomes constrained in the active site for hydride transfer^22,23^. Our previous studies in solution revealed that the fluorescence lifetimes of bound NADH and NADPH are sensitive to these conformations^18^. The ∼400ps lifetime of free NADH increased to 1340(±40)ps in the open conformation and 3200(±200)ps in substrate-free closed conformations. The increases for NADPH were larger, from ∼400ps to 1590(±50)ps and 4400(±200)ps respectively. The addition of reduced substrates (e.g. lactate for lactate dehydrogenase, isocitrate for isocitrate dehydrogenase) further increased each lifetime, to a maximum of 5300ps for NADPH in the closed configuration. No redox transfer can occur in these abortive ternary complexes as both cofactor and substrate are reduced. Catalytically productive ternary complexes with oxidized substrates were not, however, characterised at that time.

We have performed trFAIM to identify the range of NAD(P)H binding configurations present in living cells. This technique records a separate fluorescence decay for emission polarised parallel and perpendicular to that of the excitation light^24^. The application of appropriate fitting models then reveals both the fluorescence lifetime and rotational correlation time (inversely proportional to the rotational diffusion coefficient) for each species present in a heterogeneous population^18,25^. As pixel-by-pixel fitting of NAD(P)H fluorescence decays in live cells can typically resolve only two decay components, we increased signal to noise by summing emission from mitochondrial, cytosolic, and nuclear pixels into three separate decay curves per image. These were considered separately as their clearly contrasting NAD(P)H fluorescence intensities may imply distinct metabolic profiles. This approach revealed five decay components in HEK293 cells (Figure 1 and Supplementary Information Table S1). Three were consistent across all the subcellular compartments (between 175-186ps, 530-535ps and 1500-1779ps) but the longest lifetime varied, at 5400-5480ps in mitochondria and cytosol and 4582ps in the nucleus. As our 80MHz excitation terminates decay curve measurements at 12.5ns, reducing sensitivity to the slowest lifetime components, this likely represented a weighted average of longer lifetime species whose relative amplitudes changed between compartments. At short timescales, the trFAIM system provided fivefold improved resolution compared to our standard FLIM apparatus^16,26–29^, allowing detection of a previously unobserved species with a 24-25ps fluorescence lifetime.

**Figure 1:**
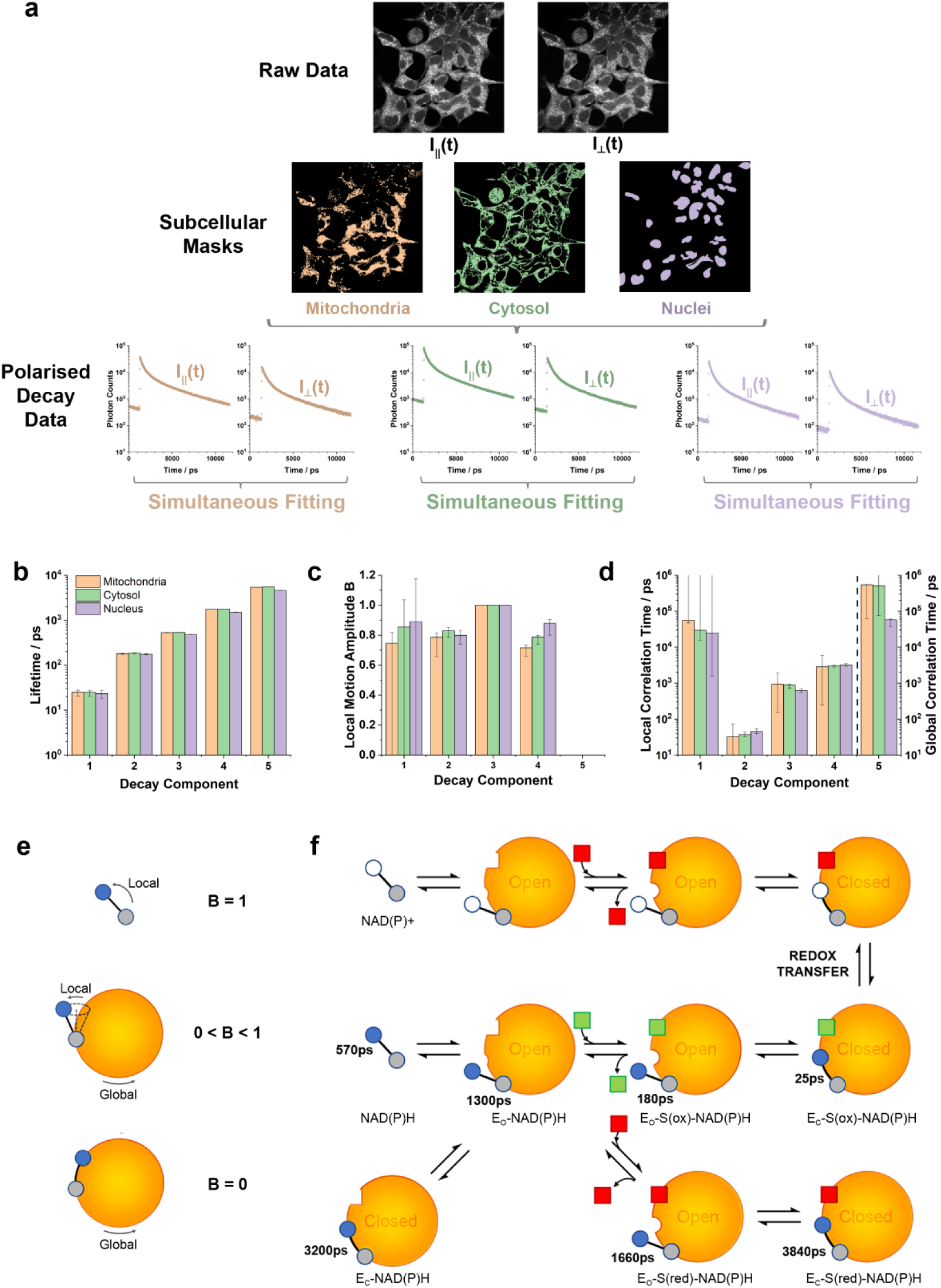
Time-resolved fluorescence anisotropy imaging (trFAIM) of NAD(P)H in HEK293 cells. (a) Masks of mitochondrial, cytosolic, and nuclear regions, generated based on their contrasting fluorescence intensities. Orthogonally polarised decay datasets were extracted by summing all photons in each image belonging to each subcellular region. (b) The fluorescence lifetimes of the five distinct species resolved and (c&d) their fluorescence anisotropy decay parameters. (e) Schematic diagram indicating the relationship between the B parameter for each component and the degree of local motion. (f) The generalised oxidoreductase reaction mechanism and the estimated fluorescence lifetimes of NADH in key binding configurations.

We next determined rotational correlation times for each of the five species identified. Based on the Stokes-Einstein-Debye equation^30^, proteins that bind NAD(P)H (10^2^-10^4^ kDa^31^) should have rotational correlation times of 30ns to 3µs (assuming an aqueous environment and a typical 1.35 g cm^-3^ protein density^32^). The longest lifetime NAD(P)H species showed rotational correlation times of 46[43,52]ns and 98[45,99]ns in the cytosol and nucleus (where square brackets indicate standard deviation confidence intervals, see Supplementary Information Table S2), consistent with enzyme binding. This increased to 689[53,691]ns in the mitochondria. While the uncertainties were significant, this large value could potentially reflect large NAD(P)H-binding complexes within this organelle, such as pyruvate dehydrogenase and complex I of the electron transport chain. Our trFAIM experiments found rotational correlation times for the 480-530ps lifetime component of 440[414,462]ps (cytosol), 450[418,477]ps (nucleus) and 502[465,534]ps (mitochondria), consistent with free NAD(P)H in aqueous solution^33^. The components with 175-186ps and 1501-1779ps lifetimes, however, showed rotational correlation times inconsistent with fully free or bound NAD(P)H. Rapid depolarisation (*τ*_rot_ = 25-39ps) was implied in the former, with the latter exhibiting times of 3422-5293ps. Such ambiguity likely occurred from the fitting model not accounting for small-scale local motion that can be present while part of a larger complex, such as NAD(P)H bound to an enzyme in an open conformation where the nicotinamide is unconstrained^18^. We therefore introduced the “wobbling in a cone” model to quantify this behaviour^25,34,35^, where a parameter *B* describes the level of rotational freedom (*B* = 0 for no local motion, *B* = 1 for free rotation, 0 < *B* < 1 for local motion while constrained). Introducing this significantly improved fit quality (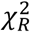 from 1.37 to 1.29, F-test P<0.002, n=18). Results showed *B* = 0.79-0.83 for the 175-186ps lifetime component and 0.79-0.88 for the 1501-1779ps component (Supplementary Information Table S3). While our fit conditions did allow for local motion in the shortest lifetime species, the slow rotational correlation time assigned to this (>25ns) instead suggested full binding. The uncertainties associated with this component were large due to its lifetime being close to the instrument resolution, and faster time-resolved fluorescence approaches will be required to probe this state in more detail. Nevertheless, the lowest values within the confidence intervals (1543-15711ps) remained longer than the rotational correlation times that would be expected for an unbound species. The assignment of the 24-25ps and 175-186ps lifetime species as enzyme-bound challenge the long-held assumption that sub-ns lifetimes in NAD(P)H FLIM purely reflect the free cofactor.

### Species in the enzyme bound population show differential responses to NAD(P) redox state

By interpreting the trFAIM results using our prior studies in solution^18^, we could assign different binding configurations to the decay components observed here inside cells. The longest lifetime (4582-5480ps) would correspond to closed, catalytically unproductive conformations with either a reduced substrate or no substrate, hereafter labelled E_C_-S(red)-NAD(P)H and E_C_-NAD(P)H respectively. The 1501-1779ps component represented open conformations, likely an average of binary and non-catalytic (reduced substrate) ternary complexes, E_O_-NAD(P)H and E_O_-S(red)-NAD(P)H. Additional routes of excited state decay must be available in the shorter lifetime (23-25ps and 175-186ps) bound states. These were likely ternary complexes with oxidized substrates, where fluorescence quenching by photoinduced electron transfer has previously been observed^36^. As this process is distance dependent^37^, the longer of these would represent the open form, E_O_-S(ox)-NAD(P)H, and the shorter the closed form, E_C_-S(ox)-NAD(P)H, supported by the lack of local motion implied by the trFAIM data. We validated these configurational assignments by generating testable predictions through mathematical modelling (Supplementary Information Appendix S1). When parameterised using rate constants from in-solution temperature jump spectroscopy experiments^22^ (Supplementary Information Table S4), our generalised model of oxidoreductase catalysis revealed dynamic variation in the steady-state concentrations of each bound species with [NAD+]:[NADH] or [NADP+]:[NADPH] (Figure 2). Notably, the model predicted an increase in the relative abundance of shorter-lifetime E_O_-S(ox)-NAD(P)H and E_C_-S(ox)-NAD(P)H species and decrease of the longer-lifetime E_O_-S(red)-NAD(P)H and E_C_-S(red)-NAD(P)H complexes as the oxidation of the cofactor pool increased. The ratio of the open forms of these species approximated to a straight line on a log-log plot against [NAD(P)+]:[NAD(P)H].

**Figure 2:**
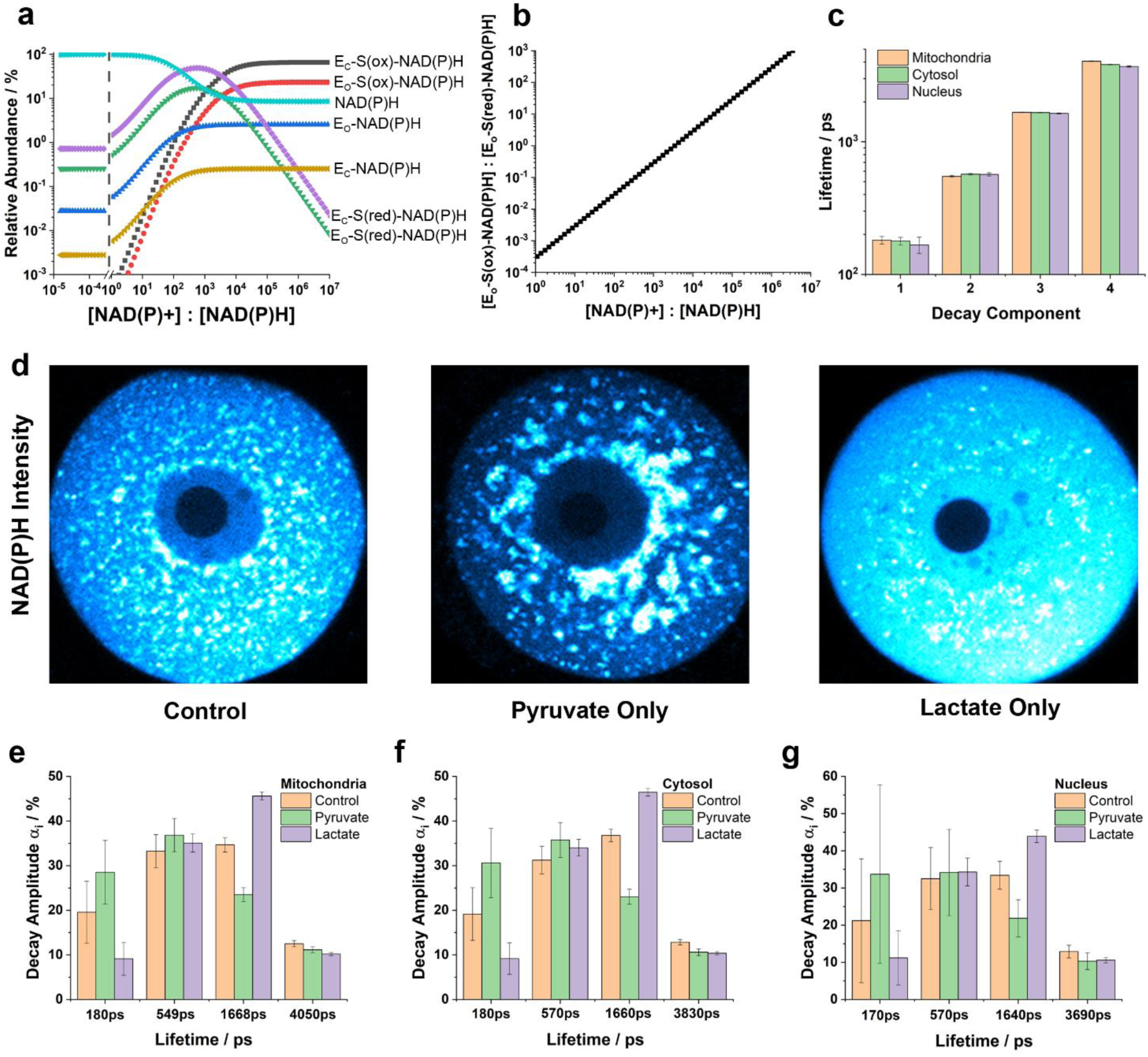
Redox-controlled conformational and binding equilibria in mammalian oocytes. (a) Variation in the relative abundance of the binding configurations with changes in [NAD+]:[NADH] (or [NADP+]:[NADPH]), predicted by a generalised model of oxidoreductase catalysis. (b) The concentration ratio of the catalytic E_O_-S(ox)-NAD(P)H to abortive E_O_-S(red)-NAD(P)H complexes followed an approximate power law relationship with the redox state. (c) This prediction was tested in standard NAD(P)H FLIM measurements on oocytes, where four fluorescence lifetimes could be resolved, ranging from 170ps to 4050ps. (d) NAD(P)H intensity images demonstrating the striking impact of media composition on the cellular NAD(P)H signal in oocytes, with lactate and pyruvate maximising and minimising the fluorescence, respectively. (e-g) Decay amplitudes in each subcellular region in response to changes in media composition.

We tested the key prediction from our model on a dataset previously acquired from studies of mammalian oocytes obtained using our standard FLIM system^16,26–29,38^. These highly specialised cells do not use glucose to produce ATP by glycolysis. Instead, exogeneous pyruvate or pyruvate derived from exogeneous lactate is imported into the mitochondria for further metabolism in the tricarboxylic acid (TCA) cycle^39^. This allows straightforward experimental control of [NAD+]:[NADH] through the choice of exogeneous substrate^40^. We compared FLIM measurements on oocytes incubated under control conditions (glucose, lactate and pyruvate) to those incubated in only lactate or only pyruvate, alongside glucose. Incubation with lactate resulted in exceptionally bright cytosolic fluorescence which became darker when the media was switched to pyruvate, reflecting the directionality of lactate dehydrogenase in each condition. Summing photon counts from all mitochondrial, cytosolic and nuclear pixels in each image initially revealed three decay components in each compartment (Supplementary Information Table S5). Under control conditions, the shortest lifetime remained constant between compartments (∼320ps), with the two longer lifetimes varying slightly (1160-1270ps, 3150-3530ps). Altered substrate supply caused variations in all three lifetimes, suggesting that there were more than three underlying fluorescent species whose relative contributions were changing. A global analysis model with lifetimes fixed between conditions, but amplitudes free to vary, extracted four components (Supplementary Information Table S6) and significantly improved fit quality (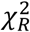 from 1.67 to 1.48, F-test P<10^-19^, n=38). The shortest three lifetimes were consistent across compartments (170-180ps, 550-570ps, 1640-1670ps), while the longest varied (4050ps in mitochondria, 3830ps in cytosol and 3690ps in the nucleus). We were unable to identify the lifetimes of individual contributors to this longest species as no further decay components could be resolved, a five-component global fit giving 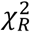 = 17.6. As in the trFAIM measurements in HEK293 cells, this likely reflected the lack of sensitivity of the experiment to longer decay times, particularly as only around 10% of the excited state population exhibited these longest lifetimes. In contrast to the trFAIM, no ∼25ps component could be resolved due to the five-fold slower time resolution of our standard FLIM instrumentation. The relative abundance of the four lifetimes was similar in each subcellular compartment. In the control media, these were 19-21%, 31-33%, 33-37% and 12-13% (shortest to longest decay components). Switching between media containing pyruvate and lactate left the amplitudes of the 550-570ps and 3690-4050ps components relatively unchanged. However, those of the other two species changed dramatically, and in opposing directions. In pyruvate-only media, the amplitude of the 170-180ps lifetime species increased to 29-34% and that of the 1640-1668ps species decreased to 22-23%. In lactate-only media, these instead decreased to 9-11% and increased to 44-46% respectively. This recreated the predictions of our model, providing support for the configurational assignment we outlined above, with the 170-180ps and 1640-1668ps decay components reflecting the E_O_-S(ox)-NAD(P)H and E_O_-S(red)-NAD(P)H species respectively.

### The redox-driven binding and conformational equilibrium impacts the biexponential decay parameters output by NAD(P)H FLIM

We next performed computational simulations of the FLIM fitting process to understand how the heterogeneous mix of binding configurations would be reflected in the simplified set of parameters (*τ*_1_, *τ*_2_ and *α*_2_) typically reported by NAD(P)H FLIM. As expected, fluorescence decays simulated with seven decay components but signal-to-noise levels reflecting pixel-by-pixel fitting were adequately described by just two components (Figure 3). With underlying lifetimes reflecting those we predicted for NADH (Supplementary Information Table S7), the shorter component (*τ*_1_) exhibited the lifetime of free NADH only at very low [NAD+]:[NADH], instead reflecting that of E_O_-S(ox)-NAD(P)H (180ps) at high ratios. It peaked between these extremes at ∼720ps due to contributions from E_O_-NAD(P)H and E_O_-S(red)-NAD(P)H, meaning *α*_2_would no longer accurately report the proportion of enzyme-bound NADH. The longer lifetime (*τ*_2_) ranged from 3500ps at low [NAD+]:[NADH] to 850ps at high ratios. The longer of these reflected the combined contribution of E_O_-S(red)-NAD(P)H and E_C_-S(red)-NAD(P)H, while the shorter value likely resulted from a mixture of E_O_-NAD(P)H and free NADH no longer assigned to *τ*_1_. With the underlying lifetimes switched to reflect those we predicted for NADPH, the behaviour of *τ*_1_ and *α*_2_ with [NADP+]:[NADPH] was broadly unchanged. *τ*_2_ was similar as for NADH at more oxidised redox states, but it reached a much larger value of around 4500ps as the cofactor pool became more reduced, due to the intrinsically longer lifetimes of many of the NADPH-associated bound species that we previously observed in solution^18^. These results demonstrated that, while the biexponential analysis typically applied to NAD(P)H FLIM data could not separately resolve the heterogeneous population of bound species underlying it, correlations with the NAD and NADP redox states nevertheless existed. To confirm these experimentally, we performed measurements on a range of multicellular systems with predictable differences in redox state.

**Figure 3:**
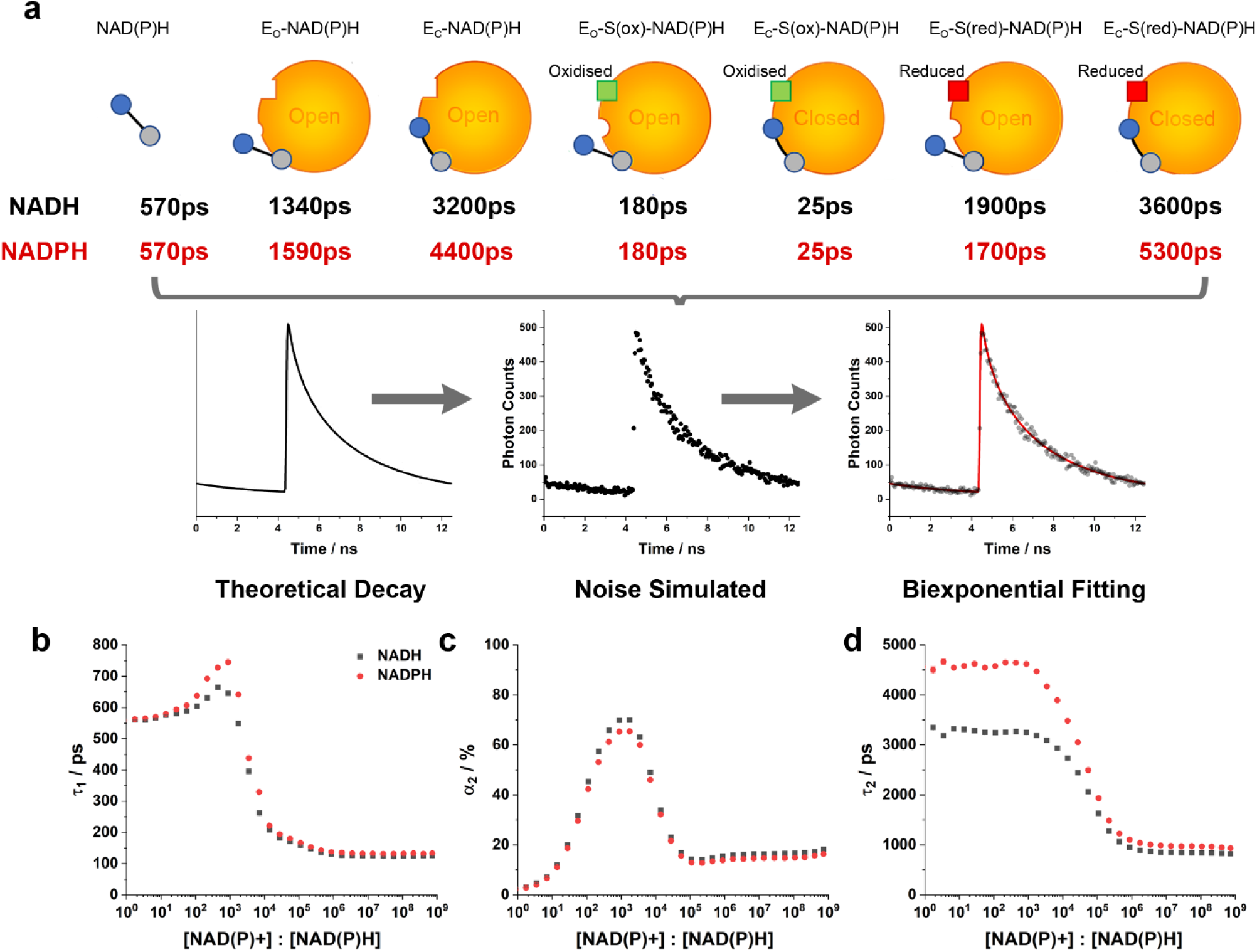
Simulating the conditions of pixel-by-pixel NAD(P)H FLIM. (a) Schematic diagrams of the generation of fluorescence decay curves based on the model-predicted values for the relative abundances of each binding configuration at varying [NAD(P)+]:[NAD(P)H]. Their lifetimes were estimated for both NADH and NADPH from the results of the present study and those obtained previously in solution^18^. 1000 iterations of noise generation and fitting were performed at varying redox ratios, allowing the canonical biexponential FLIM parameters (b) *τ*_1_, (c) *α*_2_ and (d) *τ*_2_ to be plotted.

**Figure 4:**
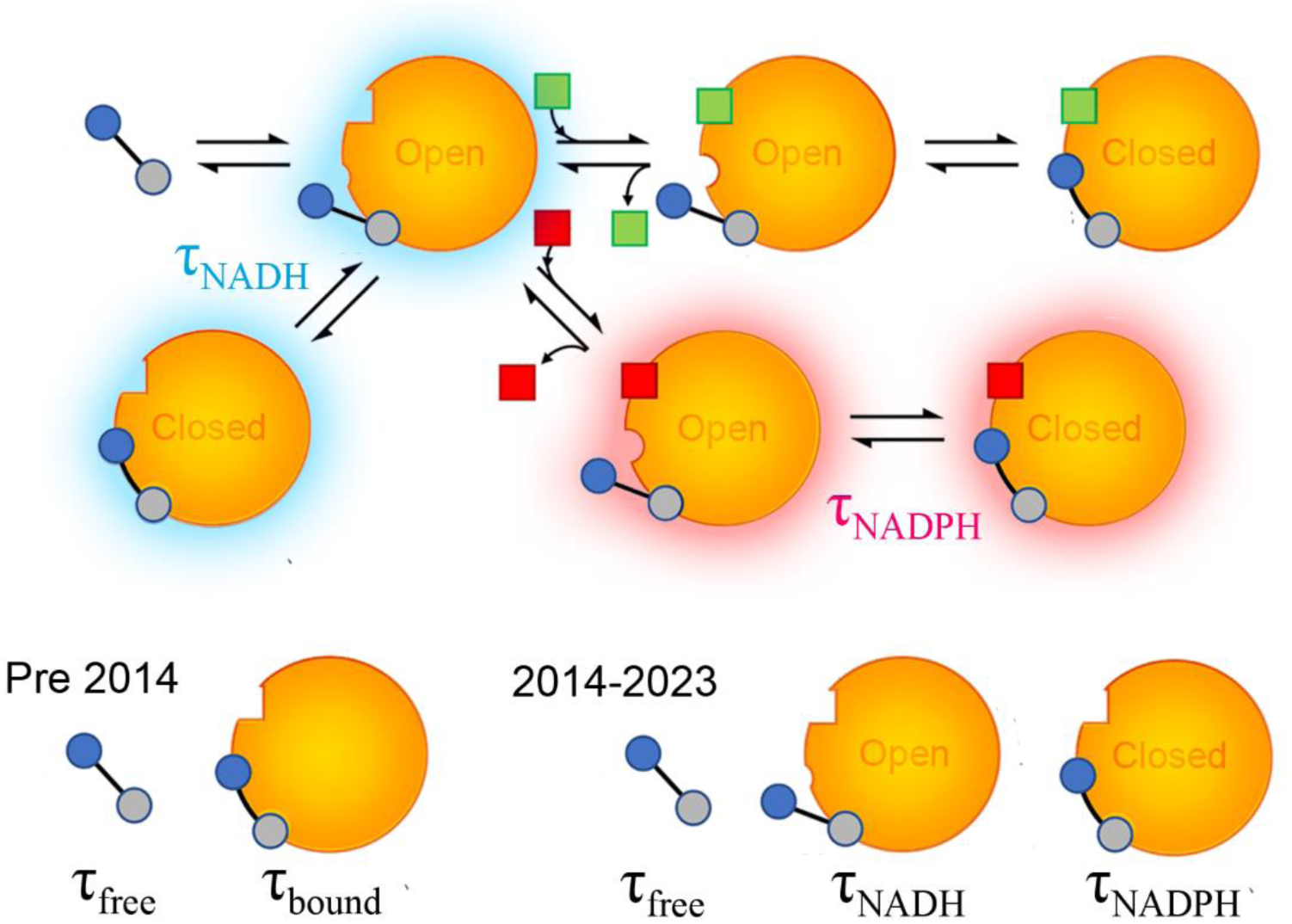
Progression in understanding the molecular basis of lifetime variations in NAD(P)H FLIM. The biexponential fluorescence decay resolved in live cell studies was originally interpreted to signify two homogeneous populations of free and bound NAD(P)H. Our prior work revealed contrasting fluorescence lifetimes of intracellular bound NADH and NADPH^16^ resulting from binding predominantly in the open and closed conformations respectively^18^. The present study provides a biochemical basis for these differences, based upon changes in the equilibrium of binding configurations driven by alterations in redox state.

In oocytes with their surrounding cumulus cells still present, *τ*_1_was significantly higher in the oocytes, at 650(±20)ps compared to 470(±20)ps (Supplementary Information Figure S1). Our modelling suggested that such large values of *τ*_1_ in the oocyte must imply a lower [NAD(P)+]:[NAD(P)H] ratio. This was in agreement with the known contrasting directionality of lactate dehydrogenase in the two cell types, with cumulus cells secreting lactate (consuming NADH) and oocytes consuming it (producing NADH)^41^. Next, in co-cultures of cortical neurons and astrocytes, we observed a longer *τ*_2_ in astrocytes compared to neurons (Supplementary Information Figure S2). Our modelling associated this with a lower [NAD+]:[NADH] in the astrocytes, a conclusion supported by equivalent measurements made with genetically encoded NAD sensors^42^. Lastly, in a mesenchymal stem cell model of oncogenesis^43^, *τ*_1_, *α*_2_ and *τ*_2_were identical in cell lines that expressed four or five oncogenes (denoted 4H and 5H), but all were significantly lower in cells expressing only three (3H, see Supplementary Information Figure S3). These differences mirrored the oxygen consumption rates of the three cell lines (Supplementary Information Table S8), with 3H respiring more slowly than 4H and 5H. As rates of respiration and redox state are closely linked^44^, this reinforced the importance of cellular redox state in controlling the fluorescence decay of NAD(P)H. However, this result also highlighted the limitations of applying a single redox equilibrium model (NAD or NADP) to interpret intracellular measurements involving both. Our model predicted that *τ*_2_should decrease with the increased [NAD+]:[NADH] expected in cells with higher respirations rates. Observation of the opposite likely reflected the greater contribution of bound NADPH, with its longer lifetime^16^, to the combined NAD(P)H signal upon NAD oxidation. Future work must model the interplay of the NAD and NADP pools, their redox states and fluorescence characteristics, but this is beyond the scope of the present study.

## Discussion

The two decay components typically resolved in NAD(P)H FLIM experiments have been widely attributed to free and enzyme-bound NAD(P)H^6,14,15^, but our study shows this to be an oversimplification. We identified enzyme-bound forms with shorter lifetimes than free NAD(P)H, specifically ternary complexes with oxidized substrates that likely accelerate non-radiative decay through photoinduced electron transfer^36^. One form had a very short lifetime (∼23-25 ps) that may fall below the time resolution of many standard FLIM systems, while another (170-180 ps) likely contributes to the faster decay component commonly assigned to free NAD(P)H. We demonstrated that the range of NAD(P)H binding configurations that contribute to its emission in living cells each exhibit distinct fluorescence lifetimes. These configurations correspond to different conformations in the catalytic action of oxidoreductase enzymes, and their relative steady state abundances are determined by the [NAD+]:[NADH] and [NADP+]:[NADPH] redox balances. This implies that the well-established correlations between NAD(P)H FLIM measurements and metabolism^6,14,15^ are underpinned by metabolically induced alterations in these ratios.

A mix of NADH and NADPH contributes to the observed signal in live cell FLIM, where we previously observed their enzyme-bound forms to possess lifetimes of ∼1500ps and ∼4400ps respectively^16^. Our results here provide a mechanism for this phenomenon. Our modelling implied that the long lifetime non-catalytic E_O_-S(red)-NAD(P)H and E_C_-S(red)-NAD(P)H species dominate when the cofactor pool is highly reduced. Such conditions are known to be maintained for the NADP pool to fulfil its role in supporting reductive biosynthesis^45^. The limiting lifetime value at low [NADP+]:[NADPH] was ∼4500ps, in striking agreement with our previous predictions for the intracellular lifetime of bound NADPH^16^. Conversely, the NAD pool is maintained in a more oxidised state to provide electron acceptors for catabolic redox reactions^44^. However, the limiting lifetime value at high [NAD+]:[NADH] here was ∼970ps. This was significantly shorter than our previous prediction for the fluorescence lifetime of bound NADH in cells, which was closely reflected by the 1340ps lifetime of the E_O_-NAD(P)H species. This shortening most probably resulted from an overestimation of the population of species with shorter lifetimes (catalytic ternary complexes and free NADH) under the most oxidised conditions. Furthermore, this limiting lifetime was reached at much higher [NAD+]:[NADH] ratios (>10^7^) than expected (∼10^2^) inside the cell^44^. These disagreements likely resulted from our choice of rate constants, as these were estimated from the only detailed characterisation available, performed on a lactate dehydrogenase isoform that favours the production of NADH (and pyruvate)^22^. Enzymes with rate constants that favour NAD+ production must instead dominate inside the cell to maintain the pool in an oxidised state.

The model we developed here successfully fulfilled its originally intended purpose of aiding the assignment of each fluorescent NAD(P)H species to the various oxidoreductase binding configurations. A more general model based upon the individual redox states of the NAD and NADP pools, possibly allowing them to be deduced from an NAD(P)H fluorescence decay measurement, would require representation of the diverse range of NADH and NADPH associated enzymes present in the cell. Although not all oxidoreductases involve the reversible binding of a substrate, the generalised reaction mechanism upon which we based our model could still be utilised through an appropriate choice of rate constants. For example, in complex I of the respiratory chain, the redox partner of NADH is a prosthetic flavin group, so the rate of substrate unbinding would be zero. A similar approach could account for the existence of oxidoreductases that do not utilise the open-closed conformational transition. These are, however, hypothetical, as this is widely observed across oxidoreductases^46–51^, including complex I^52^.

Parameterising a model linking NAD(P)H FLIM to the cellular NAD and NADP redox states would require quantitative characterisation of the conformational kinetics of a range of enzymes beyond those thus far performed on lactate dehydrogenase alone^22^. This also raises the question of how such a large array of different enzymes could be analytically represented. Yang et al. have made advances in this direction^53^, developing a simplified model that allowed flux through the mitochondrial respiratory chain to be predicted based on canonical biexponential NAD(P)H FLIM readouts. Their coarse graining procedure allocated all oxidoreductases within the cell into two categories; those that reduced NAD(P)+ to NAD(P)H, and those that oxidised NAD(P)H to NAD(P)+. With the mechanistic insights from our present study hitherto unavailable, an ansatz assumption was made that these possess two distinct fluorescence lifetimes. Our own numerical analysis of their model (Supplementary Information Appendix S2) revealed an inherent prediction that the NAD(P)H producing enzymes possess a longer fluorescence lifetime than the NAD(P)H consuming enzymes. Our results here provide evidence to support this, where we have shown that environments with high [NAD(P)+]:[NAD(P)H] ratios, which would require a higher abundance of the NAD(P)H consumers, do indeed exhibit a shorter value of *τ*_2_in a biexponential NAD(P)H FLIM experiment, and vice versa. Integrating their innovative modelling approach with the detailed molecular scale insights we provide here may be a promising avenue of future research.

In conclusion, we have shown that NAD(P)H FLIM measurements respond to metabolism via sensitivity to metabolically induced alterations in the redox states of the NAD and NADP pools. These drive the equilibrium of an array of contrasting NAD(P)H binding configurations, each contributing distinct decay kinetics to a highly heterogeneous fluorescent population. This new knowledge can now facilitate the development of accurate metabolic models, both qualitative and quantitative, for interpreting NAD(P)H FLIM.

## Methods

### HEK293 cell cultures

Frozen stocks were purchased from the American Type Culture Collection (ATCC) via LGC Standards (Teddington, UK). Cells were grown as monolayers in Dulbecco’s Modified Eagle Medium (DMEM, Thermo Fisher Scientific, Dartford, UK) containing fetal bovine serum (10%), glucose (25mM), sodium pyruvate (1mM), GlutaMAX supplement (2mM) and antibiotic-antimycotic (providing 100 units ml^-1^ penicillin, 100 μg ml^-1^ of streptomycin and 0.25 μg mL^-1^ of amphotericin B) within sterile 75cm^2^ culture flasks (Nunc EasYFlask, Thermo Fisher Scientific, Dartford, UK) in a 37 °C, 5% CO_2_ incubator. Cells were plated for imaging in glass bottomed 35mm dishes (FluoroDish, World Precision Instruments, Hitchin, UK) at a density of 300,000 per dish.

### Oocyte culture

Germinal vesicle (GV) stage oocytes were recovered from the ovaries of hormone-primed 4-5 week old (C57Bl6xCBA)F1 hybrid mice as described previously^54^. Oocytes were released into embryo-tested M2 culture media (Merck Life Science, Dorset, UK) containing IBMX (200µM) to maintain meiotic arrest. Oocytes were washed three times in M2 media and repeated pipetting (using glass pipettes) was performed to remove cumulus cells. Imaging was performed in M2 media containing glucose (5.6mM) with or without sodium pyruvate (0.2mM) and sodium lactate (10mM).

### Cortical co-cultures

E17 embryos were extracted from an adult female rat, sacrificed using cervical dislocation in accordance with Home Office regulations. Heads were stored in 4°C phosphate-buffered saline (PBS). The brains were dissected, and the cortices were isolated, crushed, and incubated in papain solution (Merck, 0.4 mg ml^-1^ in PBS) at 37°C for 20 minutes. Post-incubation, the tissue was treated with DNAse (Merck, 0.4 mg ml^-1^) and triturated to dissociate the cells with an 18G blunt filter needle and syringe. The homogenised solution was centrifuged at 500g for 5 minutes, and the cell pellet was resuspended in neurobasal medium supplemented with GlutaMAX, B-27 (Thermo Fisher) and penicillin-streptomycin. Cells were plated on poly-L-lysine-treated coverslips in the wells of a 6 well plate and incubated in 5% CO_2_ at 37°C. Medium changes were performed 48 hours after plating and every 3-4 days thereafter. Imaging was performed in recording buffer containing NaCl (150mM), KCl (4.25mM), NaH_2_PO_4_ (1.25mM), NaHCO_3_ (4mM), CaCl_2_ (1.2mM), HEPES (10mM), glucose (10mM) and MgCl_2_ (1.2mM).

### Transformed mesenchymal stem cell culture

MSC lines were gifted by Dr. Juan Manuel Funes (UCL Cancer Institute). Cells labelled 3H expressed the catalytic subunit of human telomerase (hTERT) alongside the E6 and E7 human papillomavirus (HPV) oncogenes. 4H additionally expressed SV40 small T antigen, with further expression of an oncogenic allele of H-Ras (V12) producing the 5H cells. Each of the three lines^43^ were grown in Advanced DMEM (Thermo Fisher Scientific, Dartford, UK) supplemented with fetal bovine serum (10%), GlutaMAX (2 mM), penicillin (100 U ml^−1^) and streptomycin (100 μg ml^−1^). Cells were maintained as monolayers within sterile 75 cm^2^ tissue culture flasks (Nunc EasYFlask, Thermo Fisher Scientific, Dartford, UK) in a 37°C, 5% CO_2_ incubator. For imaging, cells were grown on sterile 22mm round coverslips in the wells of a 6 well plate (Nunc, Thermo Fisher Scientific, Dartford, UK), seeded at a density of 300,000 cells per well. Coverslips were held at the microscope using a custom made imaging chamber^55^ with cells bathed in HEPES-buffered, phenol red free DMEM (Thermo Fisher Scientific, Dartford, UK). Seven coverslips were imaged for each cell type, providing n=22, 23 and 25 images for the 3H, 4H and 5H cells respectively.

### Metabolic assays

Oxygen consumption rates were measured by high-resolution respirometry (Oxygraph-2k, Oroboros Instruments, Innsbruck, Austria). Cells were collected by centrifugation and resuspended in Dulbecco’s Modified Eagle Medium with HEPES (25mM) replacing sodium bicarbonate as the buffer. Cell density was counted for normalisation using a haemocytometer. After measuring the baseline (routine) respiration rate, oligomycin A (2.5µM), FCCP (2µM) and antimycin A (2.5µM) were sequentially added to obtain leak, maximally uncoupled, and non-mitochondrial oxygen consumption rates respectively^56,57^. Glycolysis was assayed using the Seahorse XF Glycolysis Stress Test (Agilent, Harwell, UK) according to manufacturer instructions. This quantified the extracellular acidification rate (ECAR) in glucose-free conditions, following glucose stimulation and upon ATP synthase inhibition with oligomycin to allow determination of the glycolytic rate and total glycolytic capacity.

### Time-resolved fluorescence anisotropy imaging

trFAIM measurements were performed on a multimodal time-resolved fluorescence microscope. This combined an 80MHz, near-infrared, femtosecond excitation source (Insight X3, Spectra Physics, Crewe, UK), DCS-120 laser scanning unit (Becker & Hickl, Berlin, Germany), inverted microscope (Axio Observer 7, Zeiss, Cambridge, UK) with high (1.4) numerical aperture objective (Plan-Apochromat 63x/1.4 Oil M27, Zeiss, Cambridge, UK), two ultrafast hybrid detectors (HPM-100-07, Becker & Hickl, Berlin, Germany) and time-correlated single photon counting (TCSPC) electronics (SPC-180NX, Becker & Hickl, Berlin, Germany). NAD(P)H fluorescence was acquired for two minutes per field of view with 720nm excitation and 440(±40)nm emission filtering and counts were histogrammed at 14.6ps time intervals. A polarising beamsplitter cube allowed images of the orthogonally polarised fluorescence signals *I*_||_ and *I*_⊥_ to be obtained simultaneously in separate detectors. Mitochondrial regions of interest were defined by thresholding in ImageJ (National Institutes of Health, Bethesa, MD USA) based on their bright, punctate NAD(P)H fluorescence, and nuclear pixels were identified based on their much dimmer signal.

### Fluorescence lifetime imaging microscopy

Standard NAD(P)H FLIM was performed on an upright two-photon microscope (Zeiss, Cambridge, UK) with a 1.0 NA 40x water-dipping objective as previously described^16,26–29^. Excitation was provided by a Ti:sapphire laser (Chameleon Ultra II, Coherent, Cambridge, UK) tuned to 720 nm. Emission events were registered through a 460(±25) nm filter by an external detector (HPM-100, Becker & Hickl) attached to a TCSPC electronics module (SPC-830, Becker & Hickl) with a histogram bin width of 48.8ps. Scanning was performed continuously for two minutes with a pixel dwell time of 1.6 μs.

In oocytes, tetramethylrhodamine methyl ester (TMRM, 25nM) was present to aid the identification of mitochondrial pixels. Its fluorescence was collected for a 10s burst using a 610(±30) nm emission filter with excitation provided at the same wavelength as NAD(P)H to avoid possible chromatic aberration. The 585(±15) nm emission spectrum of TMRM ensured its fluorescence did not contaminate the NAD(P)H images. Nuclear regions were identified from their characteristically lower NAD(P)H fluorescence.

### Decay curve fitting

Standard pixel-by-pixel fitting of NAD(P)H FLIM data was performed in SPCImage 7.4 (Becker & Hickl, Berlin, Germany). All other decays were fit in MATLAB R2019a (The Mathworks, Cambridge, UK) using the lsqnonlin() function. In both cases, the decay model was convoluted with a manufacturer provided theoretical form for the instrument response function (IRF) of the hybrid PMT detectors,

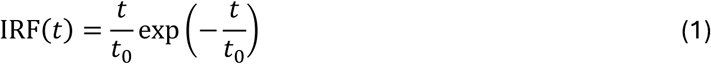

where *t*_0_, describing the width of the IRF, was included as a freely varying parameter. Under the high numerical aperture excitation and collection conditions (NA⁄*n* ∼1) utilised here, the orthogonally polarised fluorescence decays (parallel and perpendicular to the polarisation of the excitation light) were described by^58,59^,

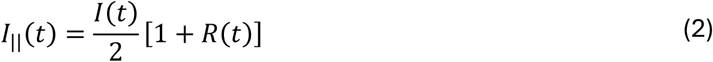

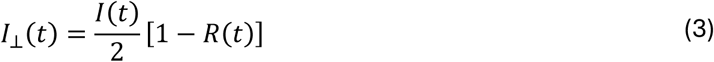

where *I*(*t*) is the intensity decay resulting from loss of the excited state population, and *R*(*t*) is the anisotropy decay resulting from diffusion-induced depolarisation. The applicability of these functional forms, including the absence of a G-factor^60^, was verified experimentally using measurements on NADH in solution (see Supplementary Information Figure S5). For trFAIM experiments, the two datasets were first summed to eliminate *R*(*t*) and the fluorescence lifetimes *τ*_*i*_ and amplitudes *α*_*i*_ were extracted by fitting a multiexponential decay function,

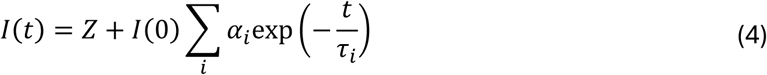

where *Z* accounted for time-uncorrelated background. The *I*(*t*) parameters were then held constant when extracting the *R*(*t*) parameters through simultaneous fitting of Equations 2 and 3. This employed the associated anisotropy function,

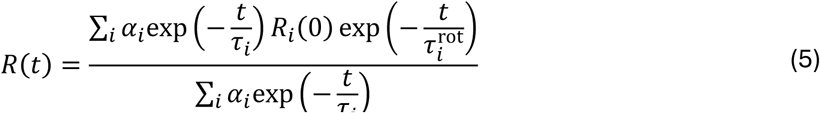

where 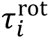 are the rotational correlation times of each species. To account for local motion, the “wobbling in a cone” model^18,25,34,35^ was later introduced,

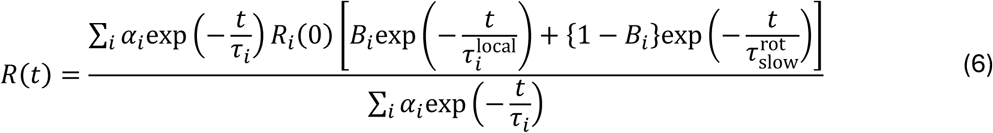

### Synthetic decay curve generation

In MATLAB R2019a (The Mathworks, Cambridge, UK), multiexponential decay functions were generated (Equation 4) with the amplitudes *α*_*i*_ provided by the solution to our modelling at a given NAD(P)+/NAD(P)H ratio. The lifetimes *τ*_*i*_ were estimated from a combination of our live-cell experiments and those performed previously in solution (see Supplementary Information Table S7). The decay was convolved with an IRF reflecting the parameters of our FLIM system (Equation 1 with *t*_0_ = 30ps, peaking at 4370ps along the time axis) and then scaled such that the peak bin had 500 counts and a constant background of 10 counts was added, typical of our FLIM measurements. The distribution of expected fit parameters at each NAD(P)+/NAD(P)H ratio was then determined by repeating, 1000 times, the addition of Poisson noise using the poissrnd() function followed by biexponential decay fitting.

### Statistical analysis

Fitting algorithms sought to minimise the reduced chi-squared function,

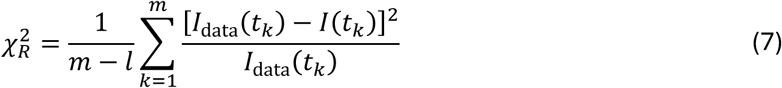

where *m* is the total number of time bins *k*, and *l* is the number of free parameters in the model. We performed statistical tests to determine whether the introduction of additional model parameters was statistically justified, evaluating the F distribution at the ratio of 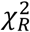 values of the two models under comparison^61^. Confidence intervals for each fitting parameter were calculated by varying them while holding all others constant until 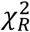 increased to a critical value determined by the F statistic at P=0.34, representing one standard deviation. Differences in FLIM parameter values between groups were assessed for statistical significance using two-tailed Student’s t-tests.

## Acknowledgements

This work was supported by Discovery Fellowship BB/W009242/1 and standard research grants BB/L020874/1 and BB/P018726/1 from the BBSRC. We thank Dr. Banyoon Cheon for preparation of the oocytes.

## Author Contributions

N.P. prepared primary cells. T.S.B., N.M., N.P., E.R.W. and M.D.E.S. performed experiments. T.S.B., E.R.W. and M.D.E.S. analysed data. T.S.B., J.C., G.S., A.J.B. and M.R.D. supervised the work. All authors drafted the manuscript.

