## Supplementary Material for "Redox-dependent binding and conformational equilibria govern the fluorescence decay of NAD(P)H in living cells"

**Table S1:** Mean NAD(P)H fluorescence lifetimes in live HEK293 cells (n=18 images) determined using trFAIM. Values in square brackets indicate average standard deviation confidence intervals.

| | **Mitochondria** | | **Cytosol** | | **Nucleus** | |
|---|---|---|---|---|---|---|
| $\tau_1$ / ps | 25 | [20, 28] | 25 | [21, 27] | 23 | [18, 28] |
| $\tau_2$ / ps | 181 | [175, 186] | 186 | [182, 190] | 175 | [169, 180] |
| $\tau_3$ / ps | 530 | [524, 535] | 531 | [527, 535] | 479 | [475, 485] |
| $\tau_4$ / ps | 1779 | [1764, 1792] | 1779 | [1767, 1790] | 1501 | [1483, 1517] |
| $\tau_5$ / ps | 5401 | [5373, 5428] | 5480 | [5453, 5507] | 4582 | [4548, 4616] |
| $a_1$ / % | 15.3 | [13.6, 15.7] | 14.9 | [13.5, 15.2] | 14.1 | [12.0, 14.4] |
| $a_2$ / % | 27.3 | [26.6, 27.8] | 29.1 | [28.6, 29.6] | 31.0 | [30.1, 31.7] |
| $a_3$ / % | 33.8 | [33.5, 34.2] | 33.7 | [33.4, 33.9] | 36.8 | [36.4, 37.2] |
| $a_4$ / % | 13.5 | [13.4, 13.7] | 12.9 | [12.8, 13.0] | 11.3 | [11.1, 11.4] |
| $a_5$ / % | 10.0 | [10.0, 10.1] | 9.4 | [9.4, 9.4] | 6.8 | [6.8, 6.9] |
| $\chi_R^2$ | 1.36 | | 1.55 | | 1.16 | |

**Table S2:** Mean associated anisotropy decay fit parameters for NAD(P)H in HEK293 cells (n=18 images).

Values in square brackets indicate average standard deviation confidence intervals.

| | **Mitochondria** | | **Cytosol** | | **Nucleus** | |
|---|---|---|---|---|---|---|
| $\tau_1^{\mathrm{rot}}$ / ps | 23420 | [92, 5e12] | 110440 | [18, 4e12] | 26198 | [11808, 4e12] |
| $\tau_2^{\mathrm{rot}}$ / ps | 32 | [25, 35] | 25 | [21, 28] | 39 | [30, 44] |
| $\tau_3^{\mathrm{rot}}$ / ps | 502 | [465, 534] | 440 | [415, 462] | 450 | [418, 477] |
| $\tau_4^{\mathrm{rot}}$ / ps | 5293 | [4031, 5562] | 3484 | [3246, 3703] | 3422 | [3258, 4031] |
| $\tau_5^{\mathrm{rot}}$ / ps | 688983 | [53149, 691367] | 46436 | [42687, 51750] | 97747 | [44517, 99042] |
| $R_1^0$ | 0.56 | [0.44, 0.57] | 0.57 | [0.46, 0.57] | 0.57 | [0.39, 0.57] |
| $R_2^0$ | 0.54 | [0.47, 0.55] | 0.55 | [0.5, 0.55] | 0.51 | [0.44, 0.53] |
| $R_3^0$ | 0.46 | [0.44, 0.48] | 0.49 | [0.48, 0.5] | 0.47 | [0.45, 0.49] |
| $R_4^0$ | 0.54 | [0.53, 0.55] | 0.56 | [0.54, 0.56] | 0.54 | [0.52, 0.54] |
| $R_5^0$ | 0.50 | [0.5, 0.51] | 0.52 | [0.51, 0.52] | 0.50 | [0.49, 0.5] |
| $\chi_R^2$ | 1.38 | | 1.56 | | 1.17 | |

**Table S3:** Mean associated "wobbling in a cone" anisotropy decay fit parameters for NAD(P)H in HEK293 cells (n=18 images). Values in square brackets indicate average standard deviation confidence intervals.

| | **Mitochondria** | | **Cytosol** | | **Nucleus** | |
|---|---|---|---|---|---|---|
| $\tau_1^{\mathrm{local}}$ / ps | 55533 | [8339, 8e12] | 29428 | [15711, 7e12] | 24759 | [1543, 8e12] |
| $\tau_2^{\mathrm{local}}$ / ps | 32 | [27, 40] | 37 | [32, 43] | 45 | [39, 53] |
| $\tau_3^{\mathrm{local}}$ / ps | 932 | [782, 983] | 883 | [726, 939] | 624 | [566, 691] |
| $\tau_4^{\mathrm{local}}$ / ps | 2852 | [2606, 3114] | 2992 | [2794, 3162] | 3201 | [2950, 3539] |
| $\tau_5^{\mathrm{slow}}$ / ps | 65783 | [50221, 72119] | 514102 | [75903, 1980698] | 57502 | [37857, 60450] |
| $R_1^0$ | 0.57 | [0.45, 0.57] | 0.57 | [0.46, 0.57] | 0.57 | [0.4, 0.57] |
| $R_2^0$ | 0.56 | [0.51, 0.56] | 0.57 | [0.53, 0.57] | 0.57 | [0.52, 0.57] |
| $R_3^0$ | 0.37 | [0.35, 0.38] | 0.39 | [0.38, 0.41] | 0.4 | [0.38, 0.42] |
| $R_4^0$ | 0.56 | [0.55, 0.57] | 0.57 | [0.55, 0.57] | 0.56 | [0.54, 0.57] |
| $R_5^0$ | 0.51 | [0.5, 0.51] | 0.5 | [0.49, 0.5] | 0.52 | [0.51, 0.52] |
| $B_1$ | 0.74 | [-974.67, 0.81] | 0.85 | [-251.54, 1.04] | 0.89 | [-132.55, 1.18] |
| $B_2$ | 0.78 | [0.72, 0.81] | 0.83 | [0.79, 0.85] | 0.81 | [0.75, 0.85] |
| $B_3$ | 1 | [fixed] | 1 | [fixed] | 1 | [fixed] |
| $B_4$ | 0.74 | [0.69, 0.76] | 0.79 | [0.74, 0.8] | 0.89 | [0.81, 0.92] |
| $B_5$ | 0 | [fixed] | 0 | [fixed] | 0 | [fixed] |
| $\chi_R^2$ | 1.32 | | 1.44 | | 1.13 | |

**Appendix S1:** Model Details

We consider a generalised oxidoreductase reaction mechanism in which redox transfer occurs in the closed enzyme conformation with both cofactor and substrate present. Both cofactor and substrate can only bind to an open enzyme, and closure is promoted by the binding of the substrate itself. This leads to the following scheme:

$$\mathrm{A} \underset{k_{-1}}{\overset{k_1}{\rightleftharpoons}} \mathrm{A{\cdot}Eo} \underset{k_{-2}}{\overset{k_2}{\rightleftharpoons}} \mathrm{A{\cdot}Eo{\cdot}S(red)} \underset{k_{-3}}{\overset{k_3}{\rightleftharpoons}} \mathrm{A{\cdot}Ec{\cdot}S(red)} \underset{k_{-4}}{\overset{k_4}{\rightleftharpoons}} \mathrm{B{\cdot}Ec{\cdot}S(ox)} \underset{k_{-5}}{\overset{k_5}{\rightleftharpoons}} \mathrm{B{\cdot}Eo{\cdot}S(ox)} \underset{k_{-6}}{\overset{k_6}{\rightleftharpoons}} \mathrm{B{\cdot}Eo} \underset{k_{-7}}{\overset{k_7}{\rightleftharpoons}} \mathrm{B}$$

where $\mathrm{A}$ is (oxidised) NAD(P)+, $\mathrm{B}$ is (reduced) NAD(P)H, $\mathrm{S}$(red) is the reduced substrate (e.g. lactate in the case of lactate dehydrogenase), $S$(ox) is the oxidised product (e.g. pyruvate) and $\mathrm{o}$ and $\mathrm{c}$ represent the open and closed conformations of the enzyme $\mathrm{E}$. We also included the possibility of "abortive" binding configurations that are not catalytically productive:

$$\mathrm{A{\cdot}Eo} \underset{k_{-i}}{\overset{k_i}{\rightleftharpoons}} \mathrm{A{\cdot}Eo{\cdot}S(ox)} \underset{k_{-j}}{\overset{k_j}{\rightleftharpoons}} \mathrm{A{\cdot}Ec{\cdot}S(ox)}$$

$$\mathrm{A{\cdot}Eo} \underset{k_{-k}}{\overset{k_k}{\rightleftharpoons}} \mathrm{A{\cdot}Ec}$$

$$\mathrm{B{\cdot}Eo} \underset{k_{-x}}{\overset{k_x}{\rightleftharpoons}} \mathrm{B{\cdot}Eo{\cdot}S(red)} \underset{k_{-y}}{\overset{k_y}{\rightleftharpoons}} \mathrm{B{\cdot}Ec{\cdot}S(red)}$$

$$\mathrm{B{\cdot}Eo} \underset{k_{-z}}{\overset{k_z}{\rightleftharpoons}} \mathrm{B{\cdot}Ec}$$

Combined, these lead to the following set of rate equations,

$$\frac{d}{dt}[\mathrm{A}] = k_{-1}[\mathrm{A{\cdot}Eo}] - k_1[\mathrm{A}][\mathrm{E}] \quad \text{(S1)}$$

$$\frac{d}{dt}[\mathrm{A{\cdot}Eo}] = k_1[\mathrm{A}][\mathrm{E}] + k_{-k}[\mathrm{A{\cdot}Ec}] + k_{-2}[\mathrm{A{\cdot}Eo{\cdot}S(red)}] + k_{-i}[\mathrm{A{\cdot}Eo{\cdot}S(ox)}] - k_k[\mathrm{A{\cdot}Eo}] - k_i[\mathrm{A{\cdot}Eo}][\mathrm{S(ox)}] - k_{-1}[\mathrm{A{\cdot}Eo}] - k_2[\mathrm{A{\cdot}Eo}][\mathrm{S(red)}] \tag{S2}$$

$$\frac{d}{dt}[\mathrm{A{\cdot}Eo{\cdot}S(red)}] = k_2[\mathrm{A{\cdot}Eo}][\mathrm{S(red)}] + k_{-3}[\mathrm{A{\cdot}Ec{\cdot}S(red)}] - k_{-2}[\mathrm{A{\cdot}Eo{\cdot}S(red)}] - k_3[\mathrm{A{\cdot}Eo{\cdot}S(red)}] \tag{S3}$$

$$\frac{d}{dt}[\mathrm{A{\cdot}Ec{\cdot}S(red)}] = k_3[\mathrm{A{\cdot}Eo{\cdot}S(red)}] + k_{-4}[\mathrm{B{\cdot}Ec{\cdot}S(ox)}] - k_{-3}[\mathrm{A{\cdot}Ec{\cdot}S(red)}] - k_4[\mathrm{A{\cdot}Ec{\cdot}S(red)}] \tag{S4}$$

$$\frac{d}{dt}[\mathrm{B{\cdot}Ec{\cdot}S(ox)}] = k_4[\mathrm{A{\cdot}Ec{\cdot}S(red)}] + k_{-5}[\mathrm{B{\cdot}Eo{\cdot}S(ox)}] - k_{-4}[\mathrm{B{\cdot}Ec{\cdot}S(ox)}] - k_5[\mathrm{B{\cdot}Ec{\cdot}S(ox)}] \tag{S5}$$

$$\frac{d}{dt}[\mathrm{B{\cdot}Eo{\cdot}S(ox)}] = k_5[\mathrm{B{\cdot}Ec{\cdot}S(ox)}] + k_{-6}[\mathrm{B{\cdot}Eo}][\mathrm{S(ox)}] - k_{-5}[\mathrm{B{\cdot}Eo{\cdot}S(ox)}] - k_6[\mathrm{B{\cdot}Eo{\cdot}S(ox)}] \tag{S6}$$

$$\frac{d}{dt}[\mathrm{B{\cdot}Eo}] = k_6[\mathrm{B{\cdot}Eo{\cdot}S(ox)}] + k_{-7}[\mathrm{B{\cdot}Eo}] + k_{-x}[\mathrm{B{\cdot}Eo{\cdot}S(red)}] + +k_{-z}[\mathrm{B{\cdot}Ec}] - k_{-6}[\mathrm{B{\cdot}Eo}][\mathrm{S(ox)}] - k_7[\mathrm{B{\cdot}Eo}] - k_x[\mathrm{B{\cdot}Eo}][\mathrm{S(red)}] - k_z[\mathrm{B{\cdot}Eo}] \tag{S7}$$

$$\frac{d}{dt}[\mathrm{B}] = k_7[\mathrm{B{\cdot}Eo}] - k_{-7}[\mathrm{B}][\mathrm{E}] \tag{S8}$$

$$\frac{d}{dt}[\mathrm{A{\cdot}Eo{\cdot}S(ox)}] = k_i[\mathrm{A{\cdot}Eo}][\mathrm{S(ox)}] + k_{-j}[\mathrm{A{\cdot}Ec{\cdot}S(ox)}] - k_{-i}[\mathrm{A{\cdot}Eo{\cdot}S(ox)}] - k_j[\mathrm{A{\cdot}Eo{\cdot}S(ox)}] \tag{S9}$$

$$\frac{d}{dt}[\mathrm{A{\cdot}Ec{\cdot}S(ox)}] = k_j[\mathrm{A{\cdot}Eo{\cdot}S(ox)}] - k_{-j}[\mathrm{A{\cdot}Ec{\cdot}S(ox)}] \tag{S10}$$

$$\frac{d}{dt}[\mathrm{A{\cdot}Ec}] = k_k[\mathrm{A{\cdot}Eo}] - k_{-k}[\mathrm{A{\cdot}Ec}] \tag{S11}$$

$$\begin{aligned}\frac{d}{dt}&[\mathrm{B{\cdot}Eo{\cdot}S(red)}] \\ &= k_x[\mathrm{B{\cdot}Eo}][\mathrm{S(red)}] + k_{-y}[\mathrm{B{\cdot}Ec{\cdot}S(red)}] - k_{-x}[\mathrm{B{\cdot}Eo{\cdot}S(red)}] \\ &- k_y[\mathrm{B{\cdot}Eo{\cdot}S(red)}]\end{aligned} \tag{S12}$$

$$\frac{d}{dt}[\mathrm{B{\cdot}Ec{\cdot}S(red)}] = k_y[\mathrm{B{\cdot}Eo{\cdot}S(red)}] - k_{-y}[\mathrm{B{\cdot}Ec{\cdot}S(red)}] \tag{S13}$$

$$\frac{d}{dt}[\mathrm{B{\cdot}Ec}] = k_z[\mathrm{B{\cdot}Eo}] - k_{-z}[\mathrm{B{\cdot}Ec}] \tag{S14}$$

We also assumed a constant concentration of available enzymes,

$$\begin{aligned}[\mathrm{E}] = [\mathrm{E_{total}}] &- [\mathrm{A{\cdot}Eo}] - [\mathrm{A{\cdot}Ec}] - [\mathrm{A{\cdot}Eo{\cdot}S(ox)}] - [\mathrm{A{\cdot}Ec{\cdot}S(ox)}] - [\mathrm{A{\cdot}Eo{\cdot}S(red)}] \\ &- [\mathrm{A{\cdot}Ec{\cdot}S(red)}] - [\mathrm{B{\cdot}Eo}] - [\mathrm{B{\cdot}Ec}] - [\mathrm{B{\cdot}Eo{\cdot}S(ox)}] - [\mathrm{B{\cdot}Ec{\cdot}S(ox)}] \\ &- [\mathrm{B{\cdot}Eo{\cdot}S(red)}] - [\mathrm{B{\cdot}Ec{\cdot}S(red)}]\end{aligned} \tag{S15}$$

The system was driven into different equilibria by controlling $R$, the ratio of total product to total substrate, whose sum was fixed at value $T$,

$$R = \frac{[\mathrm{S(ox)_{total}}]}{[\mathrm{S(red)_{total}}]} \tag{S16}$$

$$[\mathrm{T}] = [\mathrm{S(ox)_{total}}] + [\mathrm{S(red)_{total}}] \tag{S17}$$

$$[\mathrm{S(ox)}] = [\mathrm{S(ox)_{total}}] - [\mathrm{A{\cdot}Eo{\cdot}S(ox)}] - [\mathrm{A{\cdot}Ec{\cdot}S(ox)}] - [\mathrm{B{\cdot}Eo{\cdot}S(ox)}] - [\mathrm{B{\cdot}Ec{\cdot}S(ox)}] \tag{S18}$$

$$\begin{aligned}[\mathrm{S(red)}] = [\mathrm{S(red)_{total}}] &- [\mathrm{A{\cdot}Eo{\cdot}S(red)}] - [\mathrm{A{\cdot}Ec{\cdot}S(red)}] - [\mathrm{B{\cdot}Eo{\cdot}S(red)}] \\ &- [\mathrm{B{\cdot}Ec{\cdot}S(red)}]\end{aligned} \tag{S19}$$

The total concentration of cofactors was also held constant,

$$[\mathrm{N}] = [\mathrm{A}] + [\mathrm{A{\cdot}Eo}] + [\mathrm{A{\cdot}Ec}] + [\mathrm{A{\cdot}Eo{\cdot}S(ox)}] + [\mathrm{A{\cdot}Ec{\cdot}S(ox)}] + [\mathrm{A{\cdot}Eo{\cdot}S(red)}] + [\mathrm{A{\cdot}Ec{\cdot}S(red)}] + [\mathrm{B}] + [\mathrm{B{\cdot}Eo}] + [\mathrm{B{\cdot}Ec}] + [\mathrm{B{\cdot}Eo{\cdot}S(ox)}] + [\mathrm{B{\cdot}Ec{\cdot}S(ox)}] + [\mathrm{B{\cdot}Eo{\cdot}S(red)}] + [\mathrm{B{\cdot}Ec{\cdot}S(red)}] \quad \text{(S20)}$$

These equations were solved in MATLAB R2019a (The Mathworks, Cambridge, UK) using the fsolve() function and the parameter values in Table S4.

**Table S4:** Values of parameters used to solve the redox equilibrium model and their justifications

| Parameter | Value | Justification |
|---|---|---|
| $k_1$ | $6.19 \times 10^6$ M$^{-1}$ s$^{-1}$ | Measured by Zhadin et al.[1] |
| $k_{-1}$ | 559 s$^{-1}$ | |
| $k_2$ | $3.043 \times 10^7$ M$^{-1}$ s$^{-1}$ | |
| $k_{-2}$ | $1.05 \times 10^5$ s$^{-1}$ | |
| $k_3$ | 940 s$^{-1}$ | |
| $k_{-3}$ | 470 s$^{-1}$ | |
| $k_4$ | 1350 s$^{-1}$ | |
| $k_{-4}$ | 1337 s$^{-1}$ | |
| $k_5$ | 210 s$^{-1}$ | |
| $k_{-5}$ | 595 s$^{-1}$ | |
| $k_6$ | 1750 s$^{-1}$ | |
| $k_{-6}$ | $2.01 \times 10^7$ M$^{-1}$ s$^{-1}$ | |
| $k_7$ | 90 s$^{-1}$ | |
| $k_{-7}$ | $5.6 \times 10^7$ M$^{-1}$ s$^{-1}$ | |
| $k_i$ | $k_2$ | Assuming the binding and unbinding of products and substrates are equally probable |
| $k_{-i}$ | $k_{-2}$ | |
| $k_j$ | $k_3$ | Assuming the kinetics of the open/closed transition is the same with product or substrate |
| $k_{-j}$ | $k_{-3}$ | |
| $k_k$ | $k_z$ | Assuming the ligand-free open/close kinetics are the same for NAD(P)+ or NAD(P)H bound |
| $k_{-k}$ | $k_{-z}$ | |
| $k_x$ | $k_{-6}$ | Assuming the binding and unbinding of products and substrates are equally probable |
| $k_{-x}$ | $k_6$ | |
| $k_y$ | $k_{-5}$ | Assuming the kinetics of the open/closed transition is the same with product or substrate |
| $k_{-y}$ | $k_5$ | |
| $k_z$ | 41667 | Inverse of the 24μs time constant measured by Deng et al.[2] for binary complex closure. |
| $k_{-z}$ | 416667 | Derived from $k_k$ and the 10% proportion of closed binary complexes measured in Blacker et al.[3] |
| [E] | $5 \times 10^{-6}$ M | Chosen to give total bound NAD(P)H populations of ~10%, in line with values obtained using FLIM. This value is similar in magnitude to estimates published by Albe et al.[4] |
| [T] | $1 \times 10^{-3}$ M | Order of magnitude estimate from Brooks et al.[5] |
| [N] | $500 \times 10^{-6}$ M | From measurements by Zhu et al.[6] |

**Table S5:** Mean NAD(P)H fluorescence decay parameters in subcellular compartments of mammalian oocytes. Values in square brackets indicate average standard deviation confidence intervals.

| | **Mitochondria** | | **Cytosol** | | **Nucleus** | |
|---|---|---|---|---|---|---|
| Control (n=17 cells) | | | | | | |
| $\tau_1$ / ps | 0.31 | [0.3, 0.33] | 0.31 | [0.3, 0.32] | 0.32 | [0.29, 0.35] |
| $\tau_2$ / ps | 1.20 | [1.19, 1.21] | 1.16 | [1.15, 1.17] | 1.27 | [1.22, 1.29] |
| $\tau_3$ / ps | 3.41 | [3.38, 3.43] | 3.15 | [3.13, 3.17] | 3.53 | [3.42, 3.59] |
| $a_1$ / % | 0.38 | [0.37, 0.39] | 0.34 | [0.33, 0.35] | 0.39 | [0.36, 0.42] |
| $a_2$ / % | 0.41 | [0.4, 0.41] | 0.42 | [0.41, 0.42] | 0.41 | [0.4, 0.42] |
| $a_3$ / % | 0.21 | [0.21, 0.22] | 0.24 | [0.24, 0.24] | 0.20 | [0.19, 0.2] |
| $\chi_R^2$ | 1.37 | | 1.42 | | 1.09 | |
| Pyruvate Only (n=10 cells) | | | | | | |
| $\tau_1$ / ps | 0.28 | [0.27, 0.29] | 0.28 | [0.27, 0.29] | 0.26 | [0.22, 0.29] |
| $\tau_2$ / ps | 1.00 | [0.99, 1.02] | 0.98 | [0.97, 1.00] | 1.05 | [1.00, 1.10] |
| $\tau_3$ / ps | 3.31 | [3.29, 3.34] | 3.12 | [3.09, 3.15] | 3.65 | [3.41, 3.72] |
| $a_1$ / % | 0.45 | [0.44, 0.46] | 0.44 | [0.43, 0.45] | 0.46 | [0.41, 0.50] |
| $a_2$ / % | 0.36 | [0.35, 0.36] | 0.36 | [0.36, 0.37] | 0.38 | [0.36, 0.39] |
| $a_3$ / % | 0.19 | [0.19, 0.19] | 0.20 | [0.19, 0.20] | 0.16 | [0.16, 0.17] |
| $\chi_R^2$ | 1.37 | | 1.29 | | 1.06 | |
| Lactate Only (n=11 cells) | | | | | | |
| $\tau_1$ / ps | 0.39 | [0.38, 0.4] | 0.38 | [0.37, 0.39] | 0.34 | [0.32, 0.36] |
| $\tau_2$ / ps | 1.28 | [1.28, 1.29] | 1.22 | [1.21, 1.23] | 1.15 | [1.13, 1.16] |
| $\tau_3$ / ps | 3.02 | [3.01, 3.04] | 2.80 | [2.79, 2.81] | 2.77 | [2.75, 2.8] |
| $a_1$ / % | 0.33 | [0.33, 0.34] | 0.30 | [0.29, 0.31] | 0.30 | [0.29, 0.32] |
| $a_2$ / % | 0.44 | [0.44, 0.45] | 0.44 | [0.44, 0.45] | 0.43 | [0.43, 0.44] |
| $a_3$ / % | 0.22 | [0.22, 0.23] | 0.26 | [0.26, 0.26] | 0.26 | [0.26, 0.27] |
| $\chi_R^2$ | 2.22 | | 2.26 | | 1.23 | |

**Table S6:** Mean NAD(P)H fluorescence decay parameters in mammalian oocytes with compartmentalised lifetimes shared between conditions. Values in square brackets indicate average standard deviation confidence intervals.

| | **Mitochondria** | | **Cytosol** | | **Nucleus** | |
|---|---|---|---|---|---|---|
| $\tau_1$ / ps | 0.18 | [0.17, 0.19] | 0.18 | [0.17, 0.19] | 0.17 | [0.14, 0.19] |
| $\tau_2$ / ps | 0.55 | [0.54, 0.56] | 0.57 | [0.56, 0.58] | 0.57 | [0.55, 0.58] |
| $\tau_3$ / ps | 1.67 | [1.66, 1.67] | 1.66 | [1.65, 1.67] | 1.64 | [1.63, 1.65] |
| $\tau_4$ / ps | 4.05 | [4.03, 4.07] | 3.83 | [3.81, 3.85] | 3.69 | [3.65, 3.74] |
| Control (n=17 cells) | | | | | | |
| $a_1$ / % | 0.2 | [0.13, 0.26] | 0.19 | [0.13, 0.25] | 0.21 | [0.05, 0.38] |
| $a_2$ / % | 0.33 | [0.3, 0.37] | 0.31 | [0.28, 0.34] | 0.33 | [0.24, 0.41] |
| $a_3$ / % | 0.35 | [0.33, 0.36] | 0.37 | [0.35, 0.38] | 0.33 | [0.3, 0.37] |
| $a_4$ / % | 0.12 | [0.12, 0.13] | 0.13 | [0.12, 0.13] | 0.13 | [0.11, 0.15] |
| Pyruvate Only (n=10 cells) | | | | | | |
| $a_1$ / % | 0.29 | [0.21, 0.36] | 0.31 | [0.23, 0.38] | 0.34 | [0.1, 0.58] |
| $a_2$ / % | 0.37 | [0.33, 0.41] | 0.36 | [0.32, 0.4] | 0.34 | [0.23, 0.46] |
| $a_3$ / % | 0.23 | [0.22, 0.25] | 0.23 | [0.21, 0.25] | 0.22 | [0.17, 0.27] |
| $a_4$ / % | 0.11 | [0.1, 0.12] | 0.11 | [0.1, 0.11] | 0.1 | [0.08, 0.13] |
| Lactate Only (n=11 cells) | | | | | | |
| $a_1$ / % | 0.09 | [0.05, 0.13] | 0.09 | [0.06, 0.13] | 0.11 | [0.04, 0.18] |
| $a_2$ / % | 0.35 | [0.33, 0.37] | 0.34 | [0.32, 0.36] | 0.34 | [0.31, 0.38] |
| $a_3$ / % | 0.46 | [0.45, 0.46] | 0.46 | [0.46, 0.47] | 0.44 | [0.42, 0.46] |
| $a_4$ / % | 0.1 | [0.1, 0.11] | 0.1 | [0.1, 0.11] | 0.11 | [0.1, 0.11] |
| $\chi_R^2$ | 1.64 | | 1.68 | | 1.13 | |

**Table S7:** Lifetimes used in the simulation of pixel-by-pixel NAD(P)H fluorescence decays and their origin

| Species | NADH lifetime / ps | NADPH lifetime / ps | Origin |
|---|---|---|---|
| NAD(P)H | 570 | 570 | Oocyte FLIM |
| $E_O$-NAD(P)H | 1340 | 1590 | In-solution[3] |
| $E_c$-NAD(P)H | 3200 | 4400 | In-solution[3] |
| $E_O$-S(ox)-NAD(P)H | 180 | 180 | Oocyte FLIM |
| $E_c$-S(ox)-NAD(P)H | 25 | 25 | HEK293 trFAIM |
| $E_O$-S(red)-NAD(P)H | 1900 | 1700 | In-solution[3] |
| $E_C$-S(red)-NAD(P)H | 3600 | 5300 | In-solution[3] |

**Key**

| E | O | C | S(ox) | S(red) |
|---|---|---|---|---|
| Enzyme | Open | Closed | Oxidised substrate | Reduced substrate |

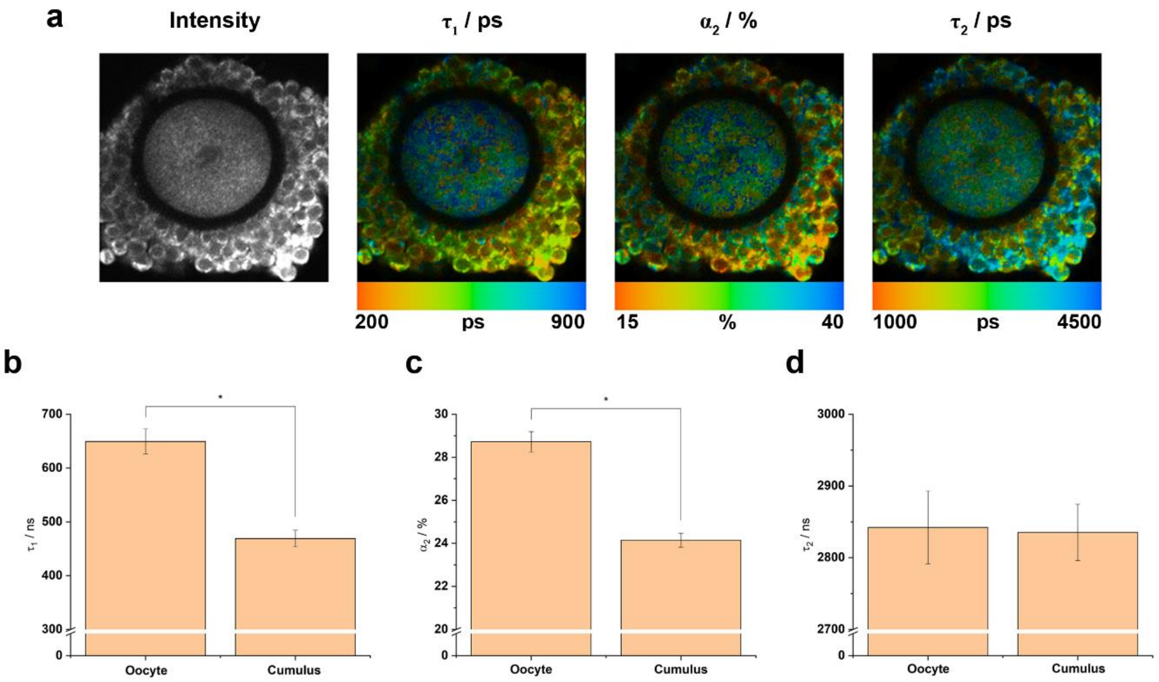


**Figure S1:** NAD(P)H FLIM in mammalian oocytes with intact cumulus cells.

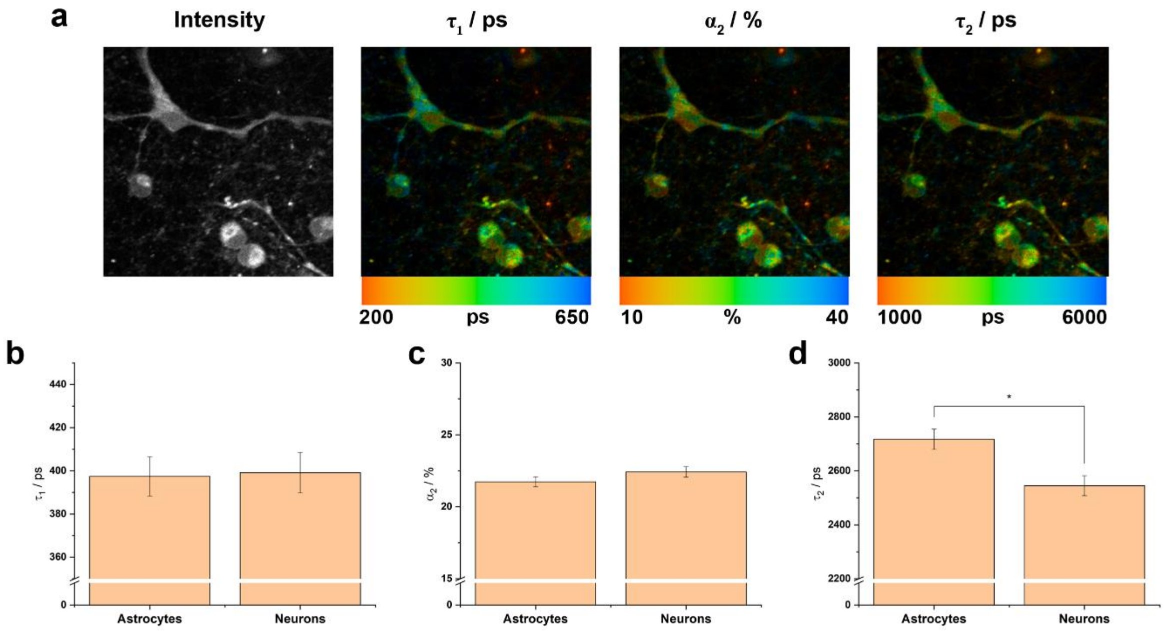


**Figure S2:** NAD(P)H FLIM in mixed co-cultures of cortical neurons and astrocytes.

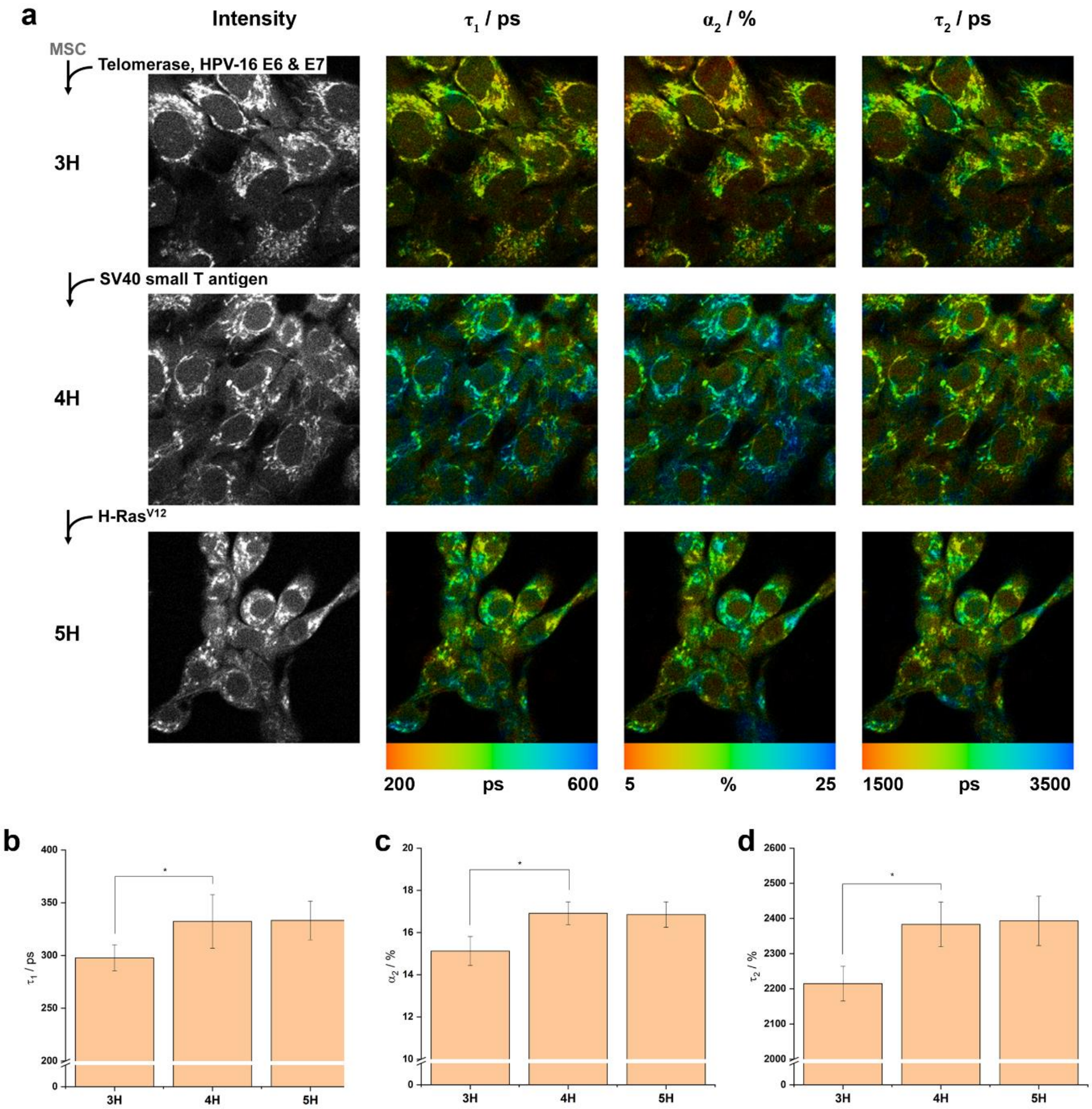


**Figure S3:** NAD(P)H FLIM in transformed mesenchymal stem cells[7].

**Table S8:** Metabolic characterisation of transformed mesenchymal stem cells.

| | **3H** | **4H** | **5H** |
|---|---|---|---|
| Oxygen Consumption / $10^{-17}$ moles $s^{-1}$ $cell^{-1}$ | | | |
| Routine | 1.6(±0.2) | 2.4(±0.5) | 2.4(±0.5) |
| Leak | 0.49(±0.06) | 0.9(±0.4) | 0.5(±0.1) |
| ETS | 2.1(±0.2) | 1.7(±0.1) | 3.3(±0.6) |
| ATP-linked | 1.1(±0.2) | 1.5(±0.2) | 1.9(±0.4) |
| Extracellular Acidification / mpH $min^{-1}$ | | | |
| Resting Glycolysis | 23(±2) | 28(±4) | 22(±1) |
| Maximum Capacity | 35(±3) | 58(±8) | 33(±2) |

**Appendix S2:** Comparisons with an existing coarse-grained model of NAD(P)H FLIM

To successfully relate NAD(P)H FLIM measurements to metabolic flux, the model developed by Yang et al. included the assumption that NAD(P)H bound to "reductases", which they define as enzymes that reduce NAD(P)+ to NAD(P)H, exhibits a different fluorescence lifetime than that bound to "oxidases", defined as enzymes that oxidise NADH to NAD+[8]. We here show that their model and calibration lead to the conclusion that the reductase lifetime $\tau_{\text{red}}$ is larger than the oxidase lifetime $\tau_{\text{ox}}$, a finding supported by the conclusions of our work.

Yang et. al define two experimentally determined parameters that relate $\tau_{\text{red}}$ and $\tau_{\text{ox}}$ to the distinct rates of NAD(P)H binding ($k_{\text{ox}}^{\text{b}}$ and $k_{\text{red}}^{\text{b}}$) and unbinding ($k_{\text{ox}}^{\text{u}}$ and $k_{\text{red}}^{\text{u}}$) to an oxidase or reductase,

$$A = (\tau_{\text{ox}} - \tau_{\text{red}}) \frac{k_{\text{ox}}^{\text{b}} + k_{\text{red}}^{\text{b}}}{k_{\text{ox}}^{\text{u}} - k_{\text{red}}^{\text{u}}} \tag{S21}$$

$$B = \frac{k_{\text{ox}}^{\text{u}} \tau_{\text{red}} - k_{\text{red}}^{\text{u}} \tau_{\text{ox}}}{k_{\text{ox}}^{\text{u}} - k_{\text{red}}^{\text{u}}} \tag{S22}$$

For convenience, we eliminate all unbinding constants using the corresponding dissociation constants,

$$A = (\tau_{\text{ox}} - \tau_{\text{red}}) \frac{k_{\text{ox}}^{\text{b}} + k_{\text{red}}^{\text{b}}}{K_D^{\text{ox}} k_{\text{ox}}^{\text{b}} - K_D^{\text{red}} k_{\text{red}}^{\text{b}}} \tag{S23}$$

$$B = \frac{K_D^{\text{ox}} k_{\text{ox}}^{\text{b}} \tau_{\text{red}} - K_D^{\text{red}} k_{\text{red}}^{\text{b}} \tau_{\text{ox}}}{K_D^{\text{ox}} k_{\text{ox}}^{\text{b}} - K_D^{\text{red}} k_{\text{red}}^{\text{b}}} \tag{S24}$$

Dividing these leads to,

$$\frac{\tau_{\text{ox}}}{\tau_{\text{red}}} = \frac{1 + \frac{k_{\text{red}}^{\text{b}}}{k_{\text{ox}}^{\text{b}}} \left(1 - \frac{A}{B} K_D^{\text{red}}\right)}{1 + \frac{k_{\text{red}}^{\text{b}}}{k_{\text{ox}}^{\text{b}}} + \frac{A}{B} K_D^{\text{ox}}} \tag{S25}$$

Yang et al. determine $A/B$ as $0.3/1.6 = 0.19$. Stinson et al. measured $K_D^{\text{red}}$~3.5μM for the "reductase" type LDH1 found in heart and $K_D^{\text{ox}}$~0.5μM for the "oxidase" type LDH5 found in muscle. Substituting these values allows us to plot the ratio of lifetimes against the ratio of binding rates (Figure S4) where it can be seen that $\tau_{\text{red}} > \tau_{\text{ox}}$.

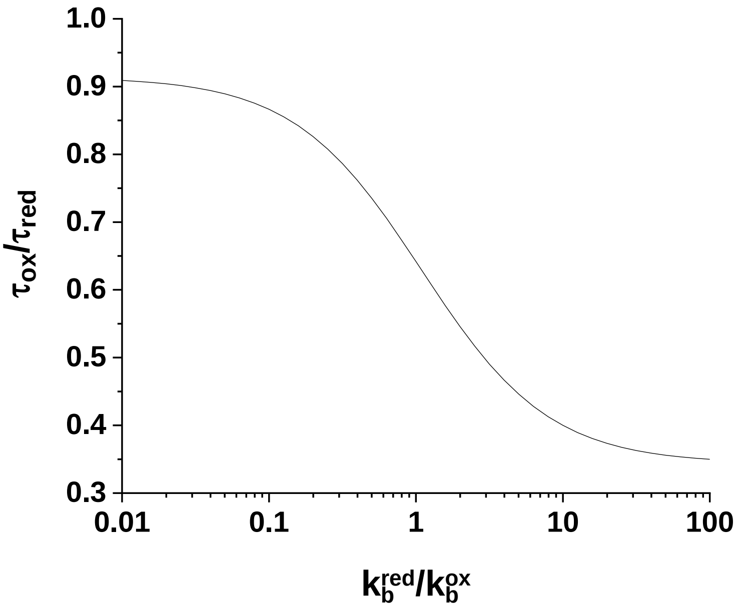


**Figure S4:** A plot of Equation S25 for experimentally determined parameter values.

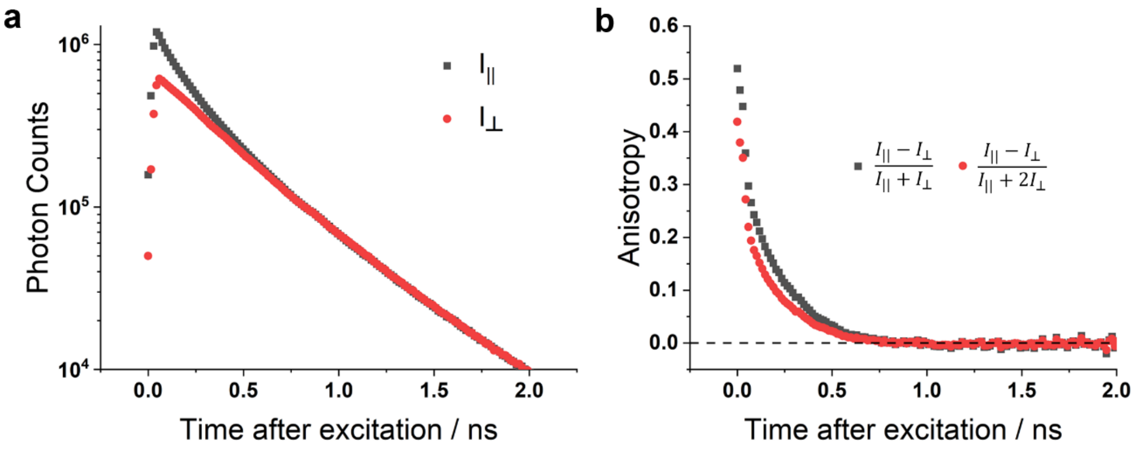


**Figure S5:** Polarised intensity decay measurements on a 1mM solution of NADH in phosphate buffered saline. (a) $I_{||}$ and $I_{\perp}$ become equal as the time after excitation increases, indicating the absence of preferential detection for either polarisation (a "G factor" of zero). (b) Appropriately formulating the intensity term of the fluorescence anisotropy for the high numerical aperture conditions of our system[9,10] leads to the expected value[11] (approximately 0.52) of the initial two-photon anisotropy for NADH in aqueous solution (black data). Ignoring these effects by assuming plane polarised excitation causes an underestimation of the anisotropy (red data).
